# Tetrapod vocal evolution: higher frequencies and faster rates of evolution in mammalian vocalizations

**DOI:** 10.1101/2023.08.09.552622

**Authors:** Matías I. Muñoz, Myriam Marsot, Jacintha Ellers, Wouter Halfwerk

## Abstract

Using the voice to produce sound is a widespread form of communication and plays an important role in contexts as diverse as parent-offspring interactions and courtship. Variation in the tempo and mode of vocal signal evolution has been studied in a phylogenetic context within orders or classes, but understanding vocal signal evolution ultimately requires comparison across all major lineages involved. Here we used comparative analyses to investigate the evolution of dominant frequency (i.e., the frequency with the highest energy content) and its association with body weight across 873 species of mammals, birds and frogs. In agreement with previous studies, we found that the negative allometric relationship between body weight and vocal frequency is a general feature of vocal systems. In addition, we found mammals to consistently vocalize at higher frequencies, and evolved their vocalizations at around 6-fold faster rates than those of birds and frogs. Although all three groups strongly rely on vocal communication, our findings show that only mammals have extensively explored the spectral acoustic space. We argue that such high vocal diversity of mammals is made possible by their unique hearing system, which evolved in the context of a small, parental-caring, nocturnal and insectivore ancestor, and has allowed them to detect, and therefore to evolve, a richer array of frequencies than other tetrapods.

## Introduction

Vocal production is widespread among animals, and communication through vocal signals occurs in numerous contexts, such as alarming, courtship and parent-offspring interactions. Besides influencing the behavior of other individuals, some animals also employ vocalizations for their own navigation and foraging, as is the case for echolocating birds and mammals (Brinkløv et al., 2013; Panyutina et al., 2017; He et al., 2021; Brinkløv et al., 2022). Vocal production thus plays a fundamental role in behavioral communication and orientation, with great potential to influence the evolutionary trajectory of lineages through effects on both survival and reproduction.

The acoustic properties of a vocalization are largely determined by the morphology of the signaler (Fitch & Hauser 2003; Ryan & Kime 2003), which sets structural limits on the sounds an animal can produce. The body size or weight of the signaler is an important attribute in dictating the frequency content of a vocalization (Martin 1971; Titze et al., 2016; García et al., 2017; García & Dunn 2019). For example, elephants are the heaviest terrestrial tetrapod alive and can communicate in the infrasonic range (i.e., frequencies lower than 20 Hz) (Herbst et al., 2012), while most ultrasound (i.e., frequencies higher than 20 kHz) producing animals weigh only a few grams (Jones 1999; Fernandes-Vargas et al. 2021). This association between weight and frequency is rooted in the fact that larger animals tend to have larger vocal organs which vibrate more efficiently at lower frequencies. Besides this, larger animals also usually have longer vocal tracts that emphasize lower frequencies through filtering phenomena (Fitch 1997; Fitch 2000). Therefore, when comparing vocalizations across multiple species it is important to consider the scaling associations between body size and the spectral content of those vocalizations (Ryan & Brenowitz 1985), especially when studying animals that span a wide range of body sizes.

Establishing and quantifying these evolutionary allometric rules is fundamental to identifying species, or groups of species, that deviate from the scaling expectation (Gould 1973), a process that in the context of vocal signals has been referred to as “allometric escape” (Tonini et al., 2020; Riondato et al., 2021). These deviating cases are interesting because they could be related, for example, to decoupled evolution between body size and the size of vocal structures (Bowling et al., 2017), or point towards morphological or behavioral specializations, such as having additional vocal structures (Charlton et al., 2013), fibrous masses attached to the vocal folds (Baugh et al., 2018) or descending the position of the larynx to extend the length of the vocal tracts (Fitch & Reby 2001). Prominent specializations would be necessary for a small bodied animal to produce extremely low frequencies, and *vice versa*. Independent of the mechanisms underlying such deviations, they are suggestive of strong past or present selection operating on vocal systems, and can therefore point us towards interesting examples of signal evolution.

To gain more insight into the evolutionary trajectories of tetrapod vocalizations we therefore designed a comparative phylogenetic study incorporating the three major lineages known to heavily rely on vocal communication, namely anurans, birds and mammals. We first assessed the allometric relationship between body weight and dominant frequency for 873 of these species to establish an acoustic allometry scaling equation across tetrapods, and to evaluate if different clades are or are not mere extensions of this general tetrapod pattern. Next, we studied acoustic allometry for each major group separately to evaluate whether they are characterized by different allometric grades (i.e., differences in their slopes and intercepts). Finally, we used a model-free approach to characterize and compare trends in the evolution of body weight and dominant frequency within frogs, birds and mammals. Our approach provides the first acoustic allometry study comparing trends between the three major lineages of vocal tetrapods in a phylogenetic framework, therefore allowing us to draw general conclusions about the factors driving the tempo and mode of vocal evolution in this diverse group of vertebrates.

## Methods

### Phylogeny and data collection

We used the tetrapod phylogenetic tree available from Chen & Wiens (2020); this time-calibrated phylogenetic tree includes 1799 species of reptiles, birds, amphibians and mammals. Species in this tree were sampled in proportion to the richness of each clade, and it has been previously used to study other aspects of tetrapod trait evolution (Anderson & Wiens 2017; Chen & Wiens 2020). We collected body weight (gr) and dominant frequency (Hz) data for as many as possible of the species of mammals, birds and frogs present in this tree. We did not collect data from other groups (i.e., non-avian reptiles, non-anuran amphibians) because information on their vocalizations is simply not widely available.

Body weight data of birds and mammals was collected from the literature and published data sets (see Data references). However, weight data was not widely available for frogs. Therefore, we collected body length data (i.e., snout-vent-lengths in mm) from Tonini et al., (2020), and transformed these values to weight (gr) using the size-weight allometric equations available from Santini et al., (2018). The association between body weight and body size in frogs depends on the microhabitat in which they live (Santini et al., 2018). Therefore, to improve the accuracy of our estimation, we collected frog micro-habitat data from Moen et al., (2016) and used micro-habitat specific allometric equations to compute frog body weights from their body lengths.

The dominant frequency of mammals was obtained from the literature and published datasets (see Data references). In some instances, dominant frequency was not measured or reported in the literature we searched, and therefore we collected other frequency measures such as the fundamental (i.e., the lowest harmonic of the call) or middle frequency (i.e., the middle point between the maximum and minimum frequencies). Dominant frequencies of frogs were obtained from the dataset available from Tonini et al., (2020). For birds, we downloaded high-quality bird songs from the online repository Xeno-canto (1-3 audio files per species) and measured their dominant frequency using a custom written R script (See details in Supplementary Materials). To corroborate the accuracy of our measures, we compared our dominant frequency measures to the measures obtained by Mikula et al., (2020), who used a similar approach to analyse songs of passerines. The dominant frequencies of passerines computed by us and by Mikula et al., (2020) were highly correlated (Pearson’s r = 0.79).

In total we collected body weight and dominant frequency data for 873 species of mammals (N = 103), birds (N = 404) and frogs (N = 366) that matched our phylogenetic tree and which were used in the subsequent comparative phylogenetic analyses.

### Comparative phylogenetic analyses

All the analyses were performed in R (R CRAN, version 4.1.1). Both body weight (gr) and dominant frequency (Hz) data were log10-transformed for all the analyses. We pruned the tetrapod phylogenetic tree from Chen & Wiens (2020) to match our dataset of 873 species. We adopted a two-pronged approach for our comparative analyses. First, we studied the allometric association between weight and frequency across all tetrapods in our sample (Methods section *Acoustic allometry across tetrapods*). Second, we focused on comparing allometric trends between mammals, birds and frogs (Methods section *Acoustic allometry between tetrapod groups*).

### Acoustic allometry across tetrapods

We measured the phylogenetic signal (i.e., the statistical tendency of closely related species to be phenotypically similar) of body weight and dominant frequency using the library ‘*phylosignal’* (version 0.8.5, Keck et al., 2016). We computed two commonly used metrics of phylogenetic signal, Pagel’s λ and Blomberg’s *K.* To evaluate the significance of phylogenetic signal estimates we compared them against estimates obtained from 10,000 permutations where trait values were randomly assigned across the phylogeny.

To quantify the allometric association between body weight and dominant frequency across all tetrapods we used phylogenetic generalised least squares (PGLS) regression as implemented in the library ‘*phylolm’* (version 2.6.2, Ho & Ané 2014). Dominant frequency was included as dependent variable and body weight as the only predictor. We fitted and compared various PGLS assuming different models of evolution for the variance-covariance matrix (i.e., Brownian motion, Pagel’s α, Ornstein-Uhlenbeck and Early Burst), and chose a Pagel’s λ model as the best fit based on their Akaike information criterion (AIC) (ΔAIC relative to the second best-fit model = 137.2). We computed 95% confidence intervals on the parameters of this best-fit model using 10,000 bootstrap replicates.

To evaluate the separate contributions of phylogeny and body weight in accounting for variation in dominant frequency we computed partial R^2^ (i.e., computed from the comparison of a full model against a reduced one) using the library ‘*rr2*’ (version 1.0.2, Ives 2019).

We used sensitivity analyses to evaluate if mammal, bird or frog lineages had a disproportionate influence on the parameters estimated from the PGLS fitted across all tetrapods. This analysis allows evaluation of whether these clades are mere extensions of the more general tetrapod allometric trend, or whether they deviate from it. For this we used the ‘*sensiPhy’* library (version 0.8.5, Paterno et al., 2018) to remove all the species from a given clade, and then compare the PGLS parameter estimates from this reduced model to the estimates computed from the model including all the species in the sample. Because species-rich clades may have a larger impact on parameter estimation (mammals are underrepresented in our sample relative to birds and frogs), this approach also implements randomization methods to correct for differences in sample size between lineages. We used 5,000 randomizations to test the influence of each one of the three clades on the slope of the PGLS model while accounting for differences in sample size.

We used sensitivity analyses to evaluate the robustness of the PGLS estimates to random reductions in the number of species sampled. We computed slope and intercept estimates after randomly removing 10, 20, 30, 40 and 50% of the species from the sample. We ran 1,000 PGLS simulations for each category of data reduction (i.e., we randomly removed 10% of species a thousand times).

To evaluate clade-specific deviations from the tetrapod allometric prediction, we further analysed the residuals of the PGLS model. We refer to these residual values computed from the linear relationship between weight and frequency as “*residual frequency*” from now on. Positive and negative residual frequency values are indicative of species that vocalize at higher and lower frequencies than those predicted for tetrapods of their weight, respectively. The magnitude of the residual value is indicative of how far (either above or below) a species is from the allometric expectation. We used the ‘*phytools’* library (version 0.7-90, Revell 2012) to map residual frequencies along the phylogenetic tree using phenograms (Revell 2013). These visual tools assume a Brownian motion (BM) model of evolution, where trait variation is undirected, and a function of the amount of time since divergence and the rate of trait evolution (σ^2^).

We tested whether residual frequency evolution followed different regimes between frogs, birds and mammals using the *ratebytree()* function in ‘*phytools’*(Revell et al., 2018) which implements the non-censored approach devised by O’Meara et al., (2006). We fitted Ornstein-Uhlenbeck (OU) models to test for differences in the trait optima (θ), pull strengths towards the optimum (α) and evolutionary rates (σ^2^) between the three clades (i.e., different evolutionary regimes). In this case, because we are comparing different subtrees, the trait optimum parameter (θ) corresponds to the estimated trait value at the root (a_0_), a quantity also known as phylogenetic mean. This method compares a model that estimates a single evolutionary regime for the three clades against alternative models where clades are allowed different regimes. For this we rescaled the total length of our tetrapod dividing it by 1000 (i.e., new root at 0.3507 instead of 350.7 million years) to improve numerical estimation (following Revell et al., 2018), and then pruned the subtrees corresponding to each major tetrapod lineage. We used maximum likelihood to estimate Pagel’s λ for residual frequencies along each subtree, and transformed the branch lengths using these values using the library ‘*geiger’* (version 2.0.11, Pennell et al., 2014). We fitted five different OU models where clades were allowed different combinations of evolutionary regimes, going from a single common regime (i.e., mammals = birds = frogs) to all regimes different (i.e., mammals ≠ birds ≠ frogs) and chose the best-fit model based on AIC.

### Acoustic allometry between tetrapod groups

After establishing the weight-frequency allometric association across tetrapods, we studied the allometric trends of each major tetrapod lineage separately. We used a least-squares phylogenetic ANCOVA (pANCOVA, Smaers & Rohlf 2016) to test for equality of slopes between mammals, birds and frogs. Equality of slopes is an assumption of pANCOVA and, as noted by Smaers et al., (2019), if slopes are unequal between lineages then these can already be considered to be characterized by different allometric grades. We used the package ‘*evomap’* (version 0.0.09000, Smaers & Mongle 2023) to compare the fit of a pANCOVA model including different slopes and intercepts for each clade (i.e., the interaction between body weight and phylogenetic group) against a single-slope single-intercept model (i.e., equivalent to the model used to establish the general tetrapod acoustic allometry), and against a model estimating different intercepts for each clade but a single slope (i.e., a model excluding the interaction between body weight and phylogenetic grouping). We chose the best fit model based on AIC. This approach is an appropriate statistical test for differences in the slopes and intercepts between phylogenetic groups because it is unbiased by sample size. The pANCOVA approach is based on variance partitioning, and is methodologically different from the sensitivity analyses described above, which test for the disproportionate influence of a clade on the general allometric trend by excluding them from the calculation. After establishing the best-supported model, we fitted this model using the ‘*phylolm’* package (version 2.6.2, Ho & Ané 2014) to obtain model estimates and corresponding confidence intervals computed from 10,000 bootstrap replicates.

After running the pANCOVA analysis, we pruned the subtrees of mammals, birds and frogs, and subset the corresponding body weight and dominant frequency data. We used these pruned trees and data to measure the phylogenetic signal of both traits using the Pagel’s λ and Blomberg’s K statistics, and computed partial R^2^ to measure the contribution of body weight and phylogeny in explaining variation in dominant frequency within each clade using the same methods and packages used for the complete dataset (see *Acoustic allometry across tetrapods* section).

### A model-free approach: The normalized Laplacian spectrum

Most comparative phylogenetic methods, and all the analyses presented so far in this article, rely on assuming a model of evolution (i.e., a mathematical description of how biologists think evolution occurs along a tree). Classically, Brownian motion and its extensions are used to study the evolution of traits along phylogenetic trees. Macroevolutionary tools that do not rely on such assumptions can provide complementary insights when combined with more classical model-based methods. We used the spectrum of the normalized graph Laplacian to characterize and compare patterns of trait evolution within mammals, birds and frogs (Lewitus et al., 2020). In brief, this analysis combines phylogenetic and trait distances between species to construct the so-called normalized modified graph Laplacian (nMGL) without the *a priori* definition of any evolutionary model. The shape (i.e., peak, skewness, maximum value) of the spectral density of the eigenvalues of the nMGL matrix is used to characterize the structure of trait evolution along a tree. Three macroevolutionary summary descriptors can be computed from these spectra: the splitter (*_n_λ\**), the tracer (*_n_η*), and the fragmenter (*_n_Ψ*). The splitter (*_n_λ\**) corresponds to the maximum eigenvalue of the nMGL, and reflects the tendency of trait values to cluster in two monophyletic groups (i.e., bipartiteness). The tracer (*_n_η*) corresponds to the maximum height of the spectrum and positively correlates with traditional measures of phylogenetic signal (e.g., Blomberg’s K). Finally, the fragmenter (*_n_Ψ*) corresponds to the skewness of the spectrum and reflects the tendency of traits to form discrete groups in the trait space (i.e., discreteness). We used the pruned trees and data to compute these macroevolutionary metrics, and evaluate whether body weight and dominant frequency have evolved in a similar or different fashion within mammals, birds and frogs. Although this analysis does not allow quantitative comparison of the evolution of traits between different trees (it is meant to compare evolutionary trends between different traits on a given tree), it can still be helpful to qualitatively compare trends of evolution between mammals, birds and frogs.

## Results

### Acoustic allometry across vocal tetrapods

Mammals, birds and frogs spanned a wide range of body weights (from approximately 0.15 gr to 30 tons) and dominant frequencies (from approximately 17 Hz to 131 kHz). Phylogenetic signal was strong and significant for body weight (Pagel’s λ = 1.001, P < 0.001; Blomberg’s *K* = 0.892, P < 0.001) and for dominant frequency (Pagel’s λ = 0.969, P < 0.001; Blomberg’s *K* = 0.200, P < 0.001), indicating that closely related species tend to be more phenotypically similar than expected if traits were randomly distributed across the phylogeny.

Heavier tetrapod species vocalized at lower dominant frequencies than lighter species (**Figure 1**). The tetrapod weight-frequency relationship was described by the negative power law *DF* = 5889×*BW*^-0.23^, where DF is the dominant frequency in Hertz and BW is the body weight in grams (**Table 1)**. For a tetrapod of 1 kg, for example, we predicted a dominant frequency of 1163 Hz. There was a strong phylogenetic structure in the model residuals (Pagel’s λ = 0.941), and phylogeny accounted for a larger proportion of dominant frequency variance (R^2^_pred_ = 0.6593) than body weight (R^2^ = 0.1537).

**Figure 1:**
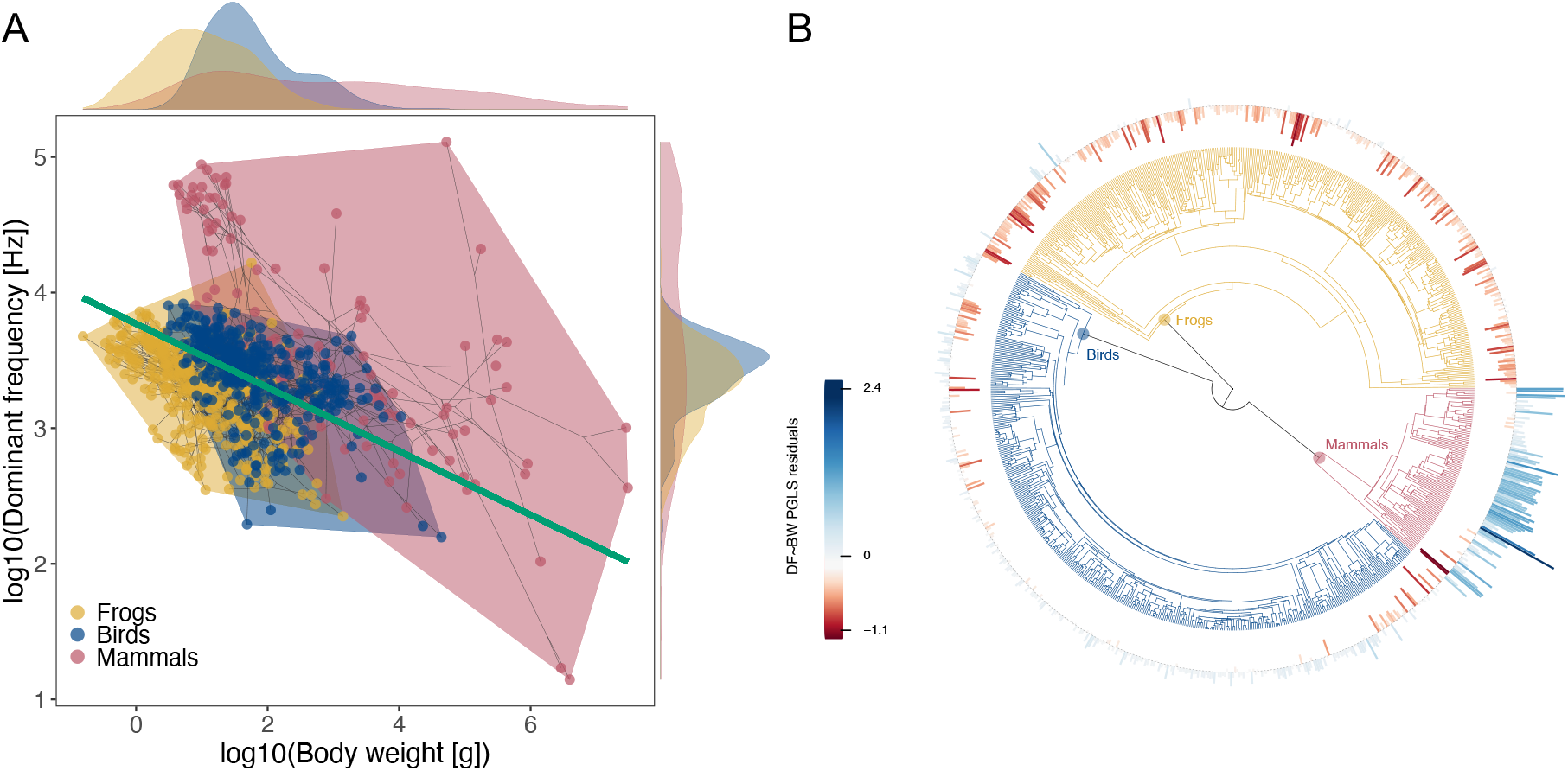
A) Phylomorphospace showing the covariation between the body weight and dominant frequency of frogs (yellow), birds (blue) and mammals (red). The green line represents the estimated phylogenetic generalized least squares (PGLS) regression computed across all species. Kernel densities are shown at the margins of each axis. B) Bar plot showing the phylogenetic residuals from the PGLS model shown in panel A) plotted at the tips of the phylogeny. Positive values are shown in blue and negative values are shown in red.

**Table 1:** Results of the PGLS model fitted across all mammal, bird and frog species in the sample.

|  | Estimate | 95% CI | S.E. | t value | P value |
| --- | --- | --- | --- | --- | --- |
| Intercept | 3.770 | [3.235, 4.280] | 0.275 | 13.695 | <0.0001 |
| log10(body weight) | -0.235 | [-0.266, -0.204] | 0.016 | -14.793 | <0.0001 |
| Pagel's $\lambda$ | 0.941 | [0.912, 0.961] | | | |
| $\sigma^2$ | 0.0011 | [0.0009, 0.0012] | | | |

95% confidence intervals were computed from 10,000 bootstrap permutations.

The tetrapod acoustic allometry was robust to reductions in sample size. Randomly removing half of the species from the analysis, for example, results only in a 1.3% average intercept change and a 7.1% average change in the slope (**Supplementary Table 1**). Sensitivity analyses also showed that mammals, birds and frogs are not mere extensions of the general tetrapod acoustic allometry. Removing mammals or birds had a substantial impact on the estimated slope (17-21% change), even after accounting for differences in sample size between clades (**Supplementary Table 2**). On the other hand, removing frogs had a non-significant effect on the slope (4.5% change, P = 0.430), but a considerable effect on the intercept (7.2% change, P < 0.0001) (**Supplementary Table 2).** Removing mammals had a significant effect on the intercept (7.4% change, P < 0.0001), but removing birds did not (2.9% change, P = 0.052, **Supplementary Table 2).** Altogether these results indicate that our estimation of the acoustic allometry across tetrapods is reliable, and that major tetrapod clades deviate in different ways from this general tetrapod allometric trend.

The variance of mammalian residual frequency values was larger than the variance of birds and frogs (**Figure 2)**. Mammals also exhibited predominantly positive residuals, indicating they vocalize at higher frequencies than expected for tetrapods of their body weight (**Figure 2**). In contrast, bird residual frequencies were mostly centred around zero, and most frogs presented negative values **(Figure 2)**. In macroevolutionary terms, these patterns suggest that the frequencies of mammal, bird and frog vocalizations have evolved at different rates towards different trait optima (i.e., under different regimes). Indeed, we found that a model that estimated a unique evolutionary regime for mammals, and a common regime for birds and frogs was best supported by the data, and significantly better than a common-regime model (**Supplementary Table 3, 4**). This best-fit model estimated a 6.1-times faster evolutionary rate for mammals (σ^2^_mammals_ = 4.352) than for birds and frogs (σ^2^_birds_ = σ^2^_frogs_ = 0.709). It also estimated a positive phylogenetic mean for mammals (a_0, mammal_ = 0.464), a mean close to zero for birds (a_0, birds_ = 0.001), and a negative one for frogs (a_0, frogs_ = –0.302). Although the pull towards the phylogenetic mean was stronger for mammals than for birds and frogs (α_mammals_ = 2.548, α_birds_ = α_frogs_ = 0.507), these values were not significantly different (*post hoc* comparison: t_100.16_ = 1.345, P = 0.182). Altogether, these results indicate that mammalian vocalizations evolved at faster rates towards frequencies higher than those allometrically predicted for tetrapods of their body weight.

**Figure 2:**
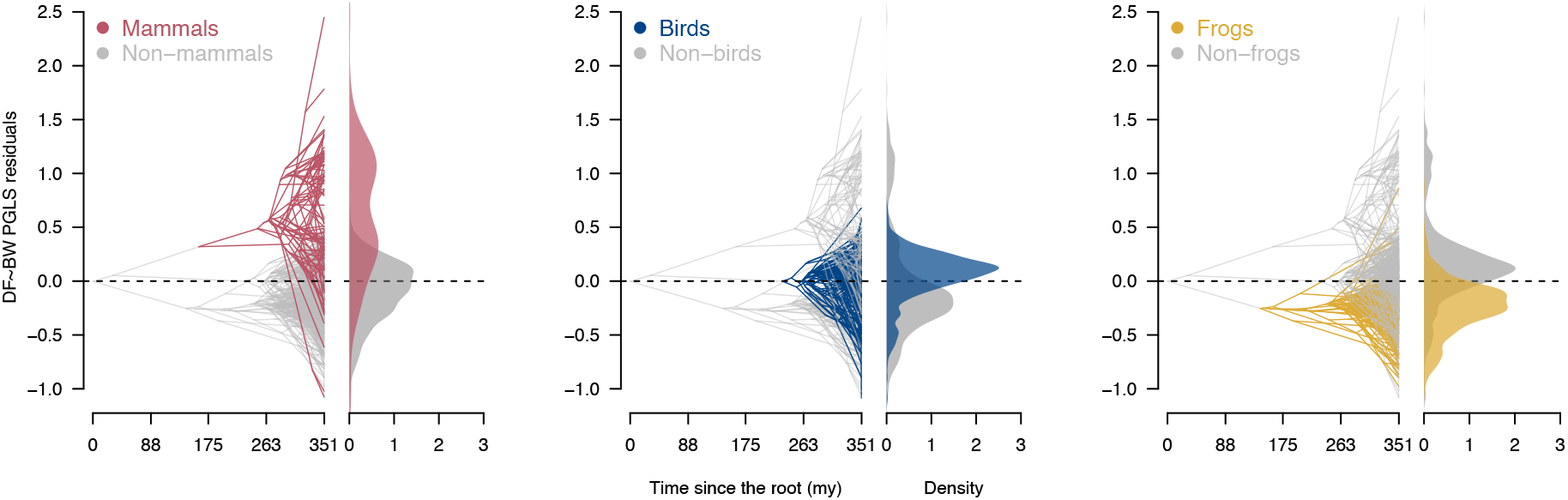
Residual frequency phenograms plotted separately for mammals (left, red), birds (centre, blue) and frogs (right, yellow). Kernel density estimates are plotted next to each phenogram, and show the distribution for each clade, as well as for those species not belonging to it (in grey). Note that the same data is contained in Fig.1B.

### Acoustic allometry within major clades of vocal tetrapods

For mammals and birds, phylogenetic signal was stronger for body weight than for dominant frequency (**Table 2**), meaning that closely related species tend to be more similar in body weight than they are in dominant frequency. This was not the case for frogs, where phylogenetic signal estimates were similar between both traits (**Table 2**).

**Table 2:** Phylogenetic signal estimated for body weight and dominant frequency data. All the phylogenetic signal estimates were significantly different from the random expectation (10,000 permutations, P < 0.0001). Note that in many cases Pagel’s λ values are at the upper theoretical limit allowed by this metric (i.e., λ = 1).

|  | Phylogenetic signal metric | Body weight | Dominant frequency |
| --- | --- | --- | --- |
| Mammals | Pagel's $\lambda$ | 1.006 | 0.966 |
|  | Blomberg's K | 1.424 | 0.758 |
| Birds | Pagel's $\lambda$ | 1.002 | 0.774 |
|  | Blomberg's K | 3.603 | 0.363 |
| Frogs | Pagel's $\lambda$ | 0.925 | 0.958 |
|  | Blomberg's K | 0.521 | 0.483 |

We found that mammals, birds and frogs are each characterized by a unique acoustic allometry grade (i.e., each clade has its own intercept and slope, **Figure 3**). A phylogenetic ANCOVA allowing different slopes and intercepts for each clade was the best-fit model (AIC = 143.31). This model provided a significantly better fit than a single-slope single-intercept model (AIC = 164.93, F_6,2_ = 7.55, P < 0.0001), and a model assuming three intercepts and a single slope (AIC = 167.45, F_6,4_ = 14.29, P < 0.0001). After accounting for variation in body size, mammals vocalized at higher mean dominant frequencies (intercept = 4.581) than birds (intercept = 3.480) and frogs (intercept = 3.442) (**Table 3**). Also, the weight-frequency association was steepest for mammals (slope = –0.344) relative to birds (slope = –0.133) and frogs (slope = –0.217) (**Table 3**). Similar to the analyses performed across all tetrapods, phylogeny explained a larger proportion of variation in dominant frequency than body weight for mammals (R^2^ phylogeny = 0.810, R^2^ body weight = 0.251), birds (R^2^ phylogeny = 0.558, R^2^_pred_ body weight = 0.034), and frogs (R^2^_pred_ phylogeny = 0.653, R^2^ body weight = 0.185). These results corroborate that mammals vocalize at higher dominant frequencies than other tetrapod lineages, and indicate that for a given evolutionary change in body weight, the associated change in dominant frequency will be larger for mammals than for birds or frogs.

**Figure 3:**
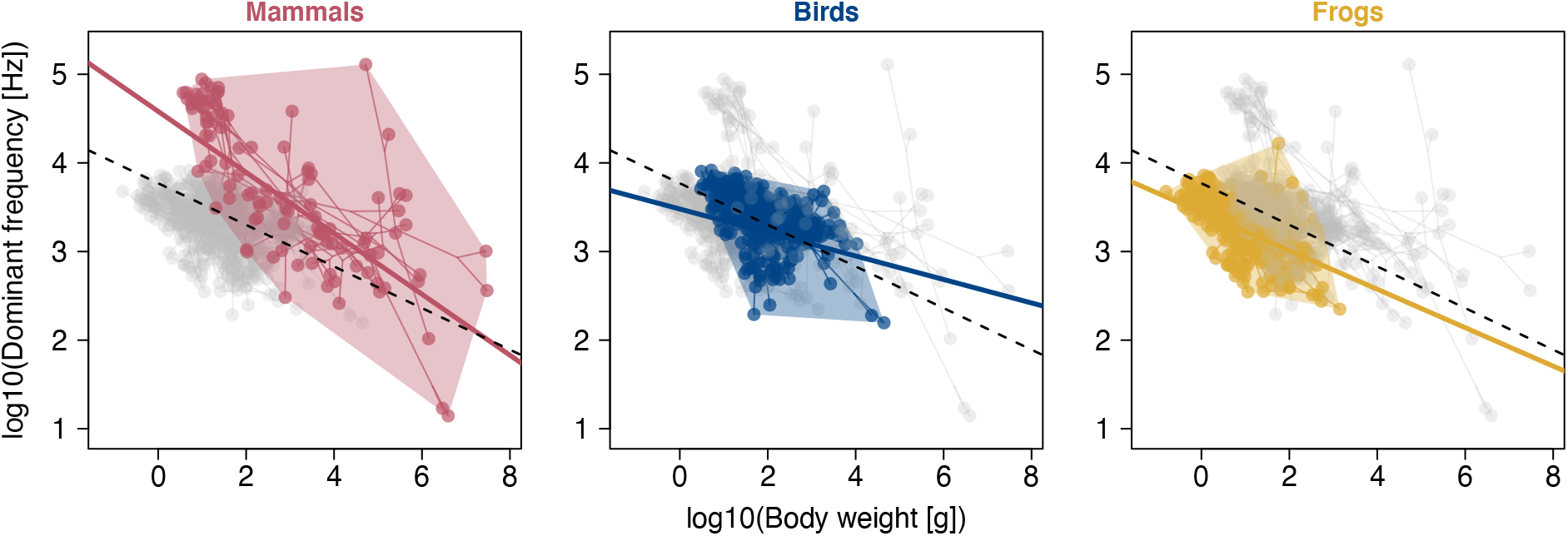
Phylomorphospace showing the covariation between the body weight and dominant frequency of mammals (red, left), birds (blue, centre) and frogs (yellow, right). The estimated PGLS line for each clade is depicted as a solid coloured line. For reference, the general tetrapod acoustic allometry (shown in Fig. 1A) is depicted as a dashed black line in each panel.

**Table 3:** Estimates of the best-fit phylogenetic ANCOVA model.

|  | Estimate | 95% CI | S.E | t value | P value |
| --- | --- | --- | --- | --- | --- |
| (Intercept) | 4.581 | [3.665, 5.476] | 0.464 | 9.869 | **<0.0001** |
| *ClassAmphibia* | -1.139 | [-2.321, 0.067] | 0.611 | -1.863 | 0.0627 |
| *ClassAves* | -1.102 | [-2.335, 0.173] | 0.631 | -1.746 | 0.0812 |
| BW | -0.344 | [-0.397, -0.291] | 0.027 | -12.791 | **<0.0001** |
| *ClassAmphibia*:BW | 0.127 | [0.057, 0.199] | 0.037 | 3.474 | **0.0005** |
| *ClassAves*:BW | 0.211 | [0.133, 0.288] | 0.040 | 5.246 | **<0.0001** |
| Pagel's $\lambda$ | 0.935 | [0.900, 0.956] | | | |
| $\sigma^2$ | 0.000987 | [0.000791, 0.001149] | | | |

95% confidence intervals were computed from 10,000 bootstrap permutations. Note that estimates and confidence intervals are given in relation to mammals.

### Normalized Laplacian spectra

Overall, the Laplacian macroevolutionary descriptors computed for body weight and body size were largely congruent (**Table 4**, **Figure 4**). For mammals, birds and frogs the values of the splitter (*_n_λ\**) and fragmenter (*_n_Ψ*) were similar for both traits (differences below 10%) (**Table 4**), indicating overall similar patterns (i.e., discreteness and bipartiteness) of body weight and dominant frequency evolution. On the other hand, for mammals and birds but not for frogs, the tracer (*_n_η*) was about 29% higher for body weight than for dominant frequency. Variation in the tracer is associated with differences in the phylogenetic signal of traits. These results corroborate the patterns observed by measuring phylogenetic signal using model-based approaches (Pagel’s λ and Blomberg’s K), and indicate that closely related mammals and birds are more similar in body weight than they are in dominant frequencies. In frogs, the Laplacian spectra of body weight and dominant frequency were strikingly similar (**Figure 4)**, indicating overall similar patterns of phenotypic evolution between these traits.

**Figure 4:**
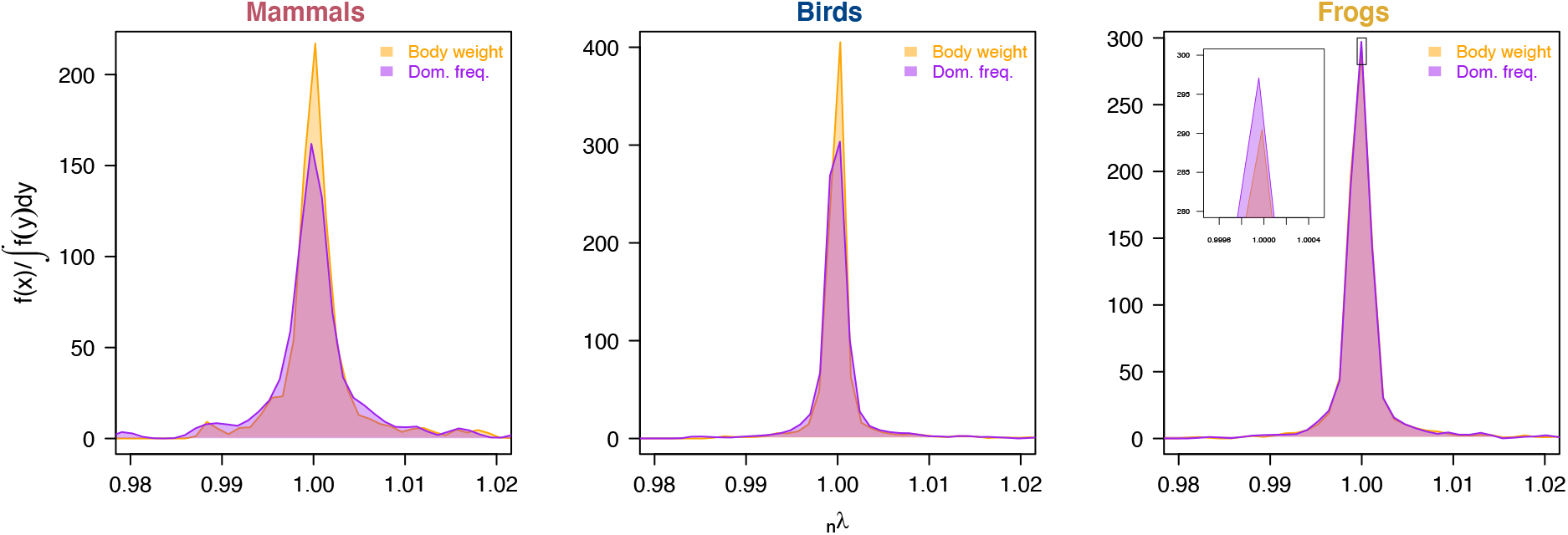
Spectral density profiles of the normalized modified graph Laplacian of body weight (orange) and dominant frequency (purple) data of mammals (right) birds (middle) and frogs (left).

**Table 4:** Macroevolutionary descriptors of the normalized Laplacian spectra computed for body weight and dominant frequency.

| Clade | Descriptor | Body weight (BW) | Dominant frequency (DF) | Difference between BW and DF (%) |
| --- | --- | --- | --- | --- |
| Mammals | Splitter (${}_n\lambda^*$) | 1.569 | 1.545 | 1.6 |
| | Tracer (${}_n\eta$) | 217.073 | 161.922 | **29.1** |
| | Fragmenter (${}_n\Psi$) | 7.955 | 7.726 | 2.9 |
| Birds | Splitter (${}_n\lambda^*$) | 1.555 | 1.507 | 3.1 |
| | Tracer (${}_n\eta$) | 405.244 | 303.418 | **28.7** |
| | Fragmenter (${}_n\Psi$) | 15.978 | 14.593 | 9.1 |
| Frogs | Splitter (${}_n\lambda^*$) | 1.499 | 1.495 | 0.3 |
| | Tracer (${}_n\eta$) | 290.462 | 297.098 | 2.3 |
| | Fragmenter (${}_n\Psi$) | 11.969 | 11.767 | 1.7 |

Differences between body weight and dominant frequency larger than 10% are highlighted in bold.

## Discussion

In their 370 million years of existence, tetrapods appear to have diversified the spectral content of their vocalizations and their body weight in a correlated fashion. Here we studied the weight-frequency allometric association in three lineages of tetrapods that rely heavily on vocal communication, spanning from frogs of less than a gram to mammals of several tons, and signals that go well beyond our human sensory capacities. We showed that the negative scaling association between weight and vocal frequency was pervasive across tetrapod lineages, although the grade of this association differs between groups. When compared to other tetrapods, we found that mammals vocalize at higher frequencies and possess more variable vocalizations, which evolved at 6-times faster rates. For mammals and birds, the phylogenetic signal of body weight was stronger than the signal of dominant frequency, meaning that closely related species within these groups are more similar in weight than they are in dominant frequency. The phylogenetic signals of frogs’ body weight and dominant frequency were virtually equal.

### Vocal systems do not explain tetrapod vocal diversity

At a proximate level, one intuitive possibility is that the vocal differences found between groups are a direct expression of differences in their vocal systems. The vocal source of mammals, birds and frogs are of different evolutionary origins, show contrasting degrees of neuromuscular control, and vocal production may even involve different cognitive processes (**Figure 5**). These differences can have important implications for both the frequency content and evolutionary diversification of their vocalizations. For example, the avian vocal organ is the syrinx, an evolutionary novelty unique to this group and located at the posterior end of the trachea (Kingsley et al., 2018). The downstream location of the avian syrinx relative to the larynx of other tetrapods improves sound production efficiency (Riede et al., 2019), and is also expected to emphasize lower frequency components due to vocal tract resonances. Despite their independent evolutionary origins and locations, the syrinx of birds and the larynx of mammals (and possibly frogs) produce sounds through a shared biomechanical mechanism (Elemans et al., 2015). Mammals and birds also share extensive neuromuscular control over their vocal organs, which can reach unprecedented degrees of detail in some birds (Adam et al., 2021). This is important, as stretching and stiffening the vocal folds (or labia in the avian syrinx) results in an increase in the frequency content of a vocalization, and can allow animals to extend their vocal frequency range beyond the limits set by the passive mass and length of their vocal organs (Titze et al., 2016). Furthermore, some groups of mammals and birds also show vocal learning (Janik & Slater 1997; Janik & Knörnschild 2021; ten Cate 2021), a cognitive capacity that could promote faster changes in the structure of their vocalizations than other evolutionary mechanisms (Rios-Chelén et al., 2012). Indeed, a recent comparative study found that vocal learning mammals deviate the most from weight-frequency scaling rules relative to non-learning ones (Garcia & Ravignani 2020). In contrast to mammals and birds, frogs are considered to have less prominent neural control over the frequency of their calls (Colafrancesco & Gridi-Papp 2016), and there is no evidence of their vocalizations being learnt (Dawson & Ryan 2009). Despite the similar biomechanical, neuromuscular and cognitive attributes of vocal production shared by many mammals and birds, only mammals have rapidly diversified the frequency of their vocalizations. This leads us to conclude that other factors not directly related to sound production might be responsible for the higher frequency and faster rates of mammalian vocal evolution.

**Figure 5:**
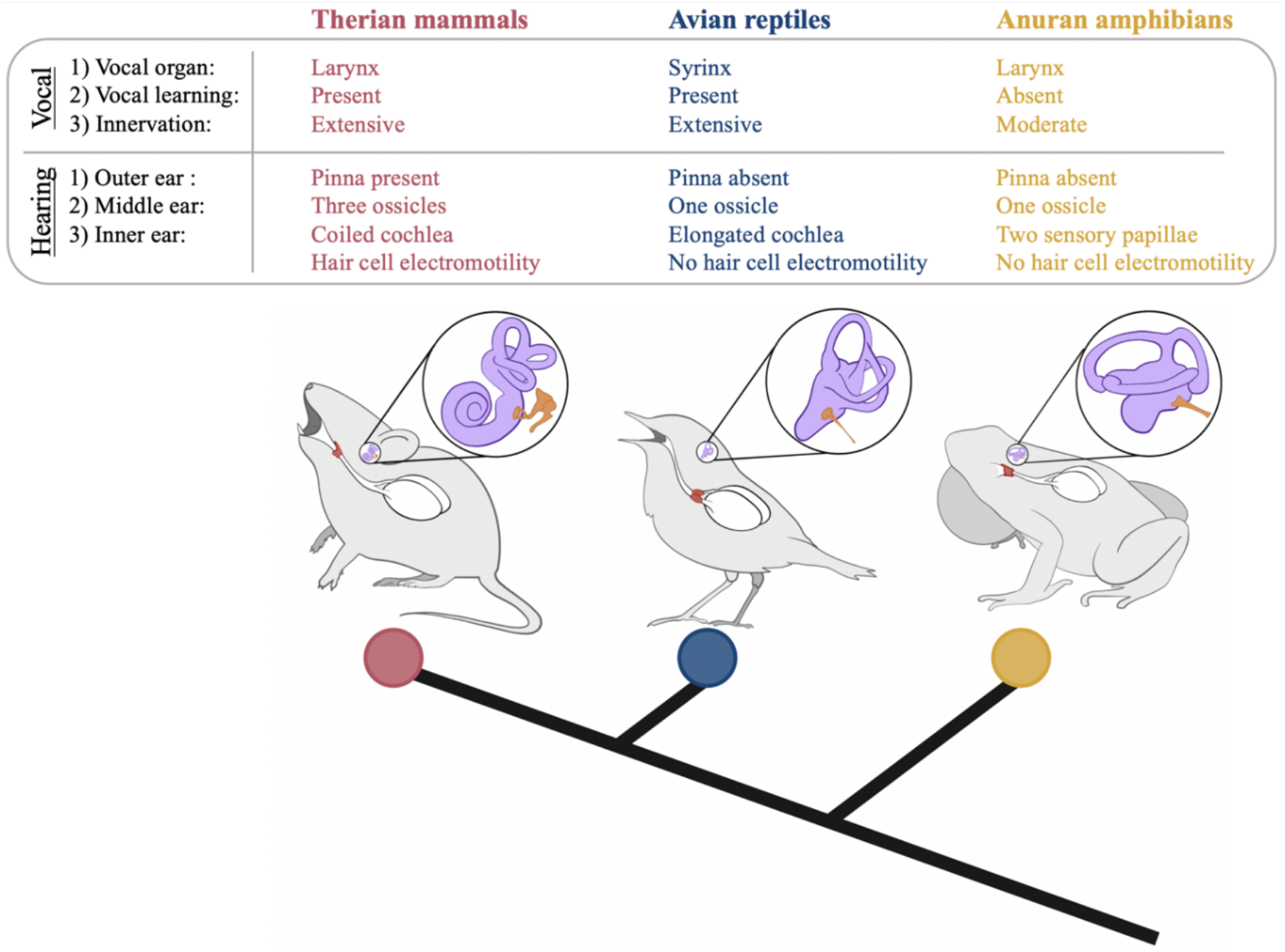
Diagram summarizing the shared and derived elements of the vocal and hearing systems of therian mammals, birds and frogs. The figure is meant to be schematic and not necessarily anatomically precise.

### Hearing systems explain tetrapod vocal diversity

> *“For an optimal combination of fascination with excellent documentation, no saga in the history of terrestrial vertebrates can match the evolution of hearing.”*
>
> — -Stephen Jay Gould (1993)

A more likely explanation for our findings is that variation in the tempo and mode of tetrapod dominant frequency evolution is indirectly permitted by differences in the auditory systems of perceivers (Endler 1992; Ryan & Cumming 2013). Following Heffner & Heffner (2008), we consider high-frequency hearing to be any noteworthy sensitivity to frequencies above 10 kHz. Mammals are well-known for their exceptional ability to hear high frequencies, often extending well-beyond 10 kHz and in some cases even reaching above 100 kHz (Masterton et al., 1969; Vater & Kössl 2011). On the other hand, with very few exceptions (see below), the hearing sensitivities of frogs and birds are typically restrained to frequencies below 10 kHz (Loftus-Hills & Johnstone 1970; Dooling et al., 2000). A handful of frog species have been described as producing and detecting calls with frequencies above 20 kHz (Feng et al., 2006; Feng & Narins 2008; Arch et al., 2009), a sensitivity limit that has not been described in birds as yet. We argue that the prominent hearing abilities of mammals have set the sensory stage that permitted the rapid diversification of their vocalizations and their shift towards higher frequencies. We present our arguments in favour of this position first from the perspective of *how* different tetrapod groups hear, and latter address *why* such outstanding hearing capacities may have evolved in mammals but not in other tetrapods.

Multiple morphological, physiological and even molecular features are responsible for the contrasting frequency sensitivities of tetrapods, and these have been extensively discussed and reviewed elsewhere (Manley 2017, 2018; Köppl 2011; Tucker 2017; Fettiplace 2020). Here we provide a brief account of the aspects that we consider to be most relevant for interpreting our results. One important difference between the hearing systems of mammals and all other tetrapods is at the level of the middle ear. The mammalian middle ear is composed of three ossicles arranged as a chain (the incus, malleus and stapes), while in the rest of the tetrapods it is composed of only the stapes (also known as columella, **Fig. 5**), which is homologous to that of mammals (Manley 2010). It is agreed that tympanic middle ears evolved independently in all major tetrapod lineages (Manley 2010; Köppl 2011; Christensen-Dalsgaard & Manley 2014; Tucker 2017). Despite their independent origins, the function of the middle ear is the same across groups: it compensates for energy losses during the transduction of air-borne mechanical vibrations at the eardrum into hydrodynamical waves at the inner ear (Christensen-Dalsgaard & Manley 2014). The single-ossicle middle ear of birds, with its cartilaginous connection to the tympanic membrane (the extra-columella), poorly transfers high frequencies into the inner ear (Manley & Gleich 1992; Gleich et al., 2004; Manley 2010), and their best hearing frequency closely matches the frequency at which the middle ear vibrates the most (Zeyl et al., 2020). This indicates that the hearing abilities of birds, and potentially also anurans with their similar middle ear configuration (Feng et al., 2006), are limited by the vibratory modes of their tympanic membranes and middle ear ossicles. In those frog species capable of ultrasonic hearing, some of the middle ear features thought to be responsible for their high-frequency sensitivities are a recessed and extremely thin tympanum, a short and light stapes, and an active mechanism to close the connection between the mouth and the middle ear cavity (Feng et al., 2006; Gridi-Papp et al., 2008). In contrast to birds and frogs that seem predominantly limited at middle ear level, it has been argued that the extended hearing range of mammals must be understood in the light of the joint contribution of both the middle and inner ears (Ruggero & Temchin 2002; Mason 2016; Köppl & Manley 2019; Luo & Manley 2020). Monotreme mammals (platypuses and echidnas), for example, possess a three-ossicle middle ear but have limited high-frequency hearing capacities compared to the rest of the mammals (the upper limit for the echidna is 16 kHz, Mills & Shepherd 2001).

Mammals are also exceptional at the inner ear level. One feature involved in expanding the hearing range of mammals is the coiled shape of their auditory inner ear organ, the cochlea (**Fig. 5**). A fully coiled cochlea is present in all therian mammals (marsupials and placentals), while in monotremes it is elongated and only slightly coiled at its apex (Schultz et al., 2017). The bird cochlea (also known as lagena) is homologous to the mammalian organ (Köppl 2011; Lipovsek & Elgoyhen 2023), but has an elongated shape and completely lacks the coiling typical of therian mammals (**Fig. 5**). Cochlear coiling allows for a longer sensory epithelium on top of which the hair cells sit (i.e., the basilar membrane), and the basilar membrane lengths of mammals (West 1985), and to some degree birds (Gleich et al., 2004), positively correlate with their hearing range. Therian mammals have long basilar membranes going from around 7 mm in the mouse to 60 mm in the elephant (West 1985). On the other hand, the basilar membranes of birds are typically shorter than 6 mm even in birds as large as the ostrich and emu (Gleich & Langemann 2011). The one avian exception is the barn owl (*Tyto alba*). Barn owls are auditory specialists that use hearing to hunt prey in the dark. They possess the longest basilar membrane described for birds (12 mm, Fisher et al., 1988), and are one of the few birds known to hear frequencies above 10 kHz, together with other owls and one species of hummingbird (Dyson et al., 1998; Duque et al., 2020). As a comparison, the basilar membranes of the cat, another nocturnal hunter, and the black rat, a nocturnal prey, are 24 and 12 mm long, respectively, and both species are capable of ultrasonic hearing (Heffner & Heffner 1985; Kelly & Masterson 1977). In contrast to the avian and mammalian conditions, the inner ear of frogs is composed of two main sensory epithelia with non-overlapping frequency selectivity, the amphibian papilla and the basilar papilla. The amphibian papilla is responsible for low-frequency hearing (<1.5 kHz), is unique to amphibians, and has a complex tonotopic organization (Lewis et al., 1982; Smotherman & Narins 2004). The anuran basilar papilla is involved in the detection of high-frequencies (up to 8.2 kHz with the exception of ultrasonic frogs, Loftus-Hill & Johnstone 1970), is relatively simple in its innervation, is not tonotopically organized, and its homology to the basilar membrane of other tetrapods is unclear (Christensen-Dalsgaard & Carr 2008; Pfaff et al., 2019). Among those few ultrasonic frogs, besides the middle ear specializations mentioned above, there are further small-scale modifications at the basilar papilla which are considered to be responsible for their ultrasonic sensitivities (Arch et al., 2012; Cobo-Cuan et al., 2020). These frog examples are outstanding, as they demonstrate that provided some adjustments are made, even a comparatively simple tympanic hearing system can allow for high-frequency hearing (see Manley & Kraus 2010 for a reptilian example).

At a molecular level, one additional factor thought to be involved in extending the auditory range of therian mammals is the presence of cochlear amplification mediated by hair cell electromotility (Ashmore 2019). The soma of mammalian outer hair cells contracts and elongates with changes in trans-membrane polarization, resulting in considerable amplification of the wave traveling along the basilar membrane. The membrane protein prestin is responsible for this somatic motility (Zheng et al., 2000; Dallos 2008), and ultrasound echolocating bats, whales and mice show convergent prestin gene sequences (Li et al., 2008; Li et al., 2010; He et al., 2021), supporting the involvement of the protein in mammalian high-frequency hearing. Birds and frogs also express a prestin homolog in their inner ears, which has been suggested to play a role in cochlear amplification in birds (Beurg et al., 2013; Fettiplace 2020). Still, these variants of the protein do not confer somatic electromotility to the hair cells expressing them, which remains an exclusive feature of the mammalian outer hair cells (He et al., 2003; Tang et al., 2013; Lipovsek & Elgoyhen 2023).

Overall, there are prominent morphological and molecular differences in *how* different groups of tetrapods hear, and these differences provide the sensory basis that permits the extended vocal diversity we found for mammals in this study. At this point it is also necessary to address *why* such different hearing sensitivities may have arisen during their particular evolutionary histories.

One possibility is that the hearing systems of modern tetrapods, together with their conserved and derived elements, are the product of historical contingencies (Christensen-Dalsgaard & Manley 2014; Manley et al., 2018). Following the thought experiment first proposed by Stephen J. Gould (1989), if we were to re-play tetrapod evolutionary history, it is possible that just by chance we would find them having different hearing systems than those we observe today. In other words, the hearing systems of modern tetrapods ended up the way they are due to contingency, and their apparent complex or simple (but adequate) organizations cannot be unequivocally attributed to predictable adaptive processes. It is also possible that at their origin, they were the product of adaptive processes not necessarily related to hearing or communication (e.g., the evolution of the mammalian middle ear is intimately linked to the evolution of their jaws, Luo & Manley 2020). Even if adaptive storytelling is an appealing approach for explaining evolution (see the critique by Gould & Lewontin 1979), historical contingency is a possibility that cannot be ruled out and deserves mentioning, especially considering the more than 350 million years of tetrapod evolutionary history and all the planetary events that happened during this timeframe.

Alternatively, it has been suggested that mammalian high-frequency hearing evolved in relation to the benefits that such a sensitivity has for sound localization (Heffner & Heffner 1992, 2008). The mammalian outer ear, the pinna, is another structure unique to mammals, and is involved in sound localization (**Fig. 5**). But, what high-frequency sounds have been relevant for mammals to detect along their evolutionary history? We can think of three possibilities.

First, it is possible that high-frequency hearing evolved in relation to communication with other conspecific individuals, and in particular with infants. Parental care is a crucial aspect of the life of all mammals, and it’s possible that their high-frequency hearing evolved in the contexts of listening to their own offspring. All three groups of living mammals produce and feed milk to their offspring, but the dependence of infants on parents is not limited to nutrition and can involve other important aspects like protection, locomotion and hygiene (Capellini & West 2016; González-Mariscal 2022). New-borns and juveniles are small, and therefore produce high-frequency sounds. It is probable that the cries of their own offspring are the highest frequency conspecific sounds that adult mammals must be able to detect and locate. In species with large litter sizes, e.g., the stem-mammal *Kayentatherium* has been associated with clutches of more than 35 perinates (Hoffman & Rowe 2018), the detection of the cries of an infant by the parents can lead to the defence of the rest of the clutch against a threat. Yet whether the hearing range of adults relates to the vocal frequency of infants has not been formally quantified, or at least not in mammals; in archosaurs (birds and crocodiles), cochlear elongation is associated with the presence of juvenile high-frequency vocalizations (Hanson et al., 2021).

High-frequency hearing could also have played a role during adult-adult communication. For a long time since the origin of the lineage, mammals and their relatives were predominantly small in body size (<100 grams) and of nocturnal habits, trends that shifted toward more diverse body sizes and patterns of diel activity somewhere around the time that dinosaurs went extinct 66 million years ago (Slater 2013; Maor et al., 2017; Benevento et al., 2023). Because of their small body size, these early mammals likely vocalized at high frequencies, although perhaps not as high as their offspring. These high-frequency vocalizations would have been useful for communicating with adult conspecifics under nocturnal low-light conditions, and would have provided a relatively private channel from communication as they would have been undetectable to crepuscular or nocturnal reptilian predators (see discussion above on the limits of non-mammalian hearing). From our results we predict dominant frequencies of 10 kHz or more for mammals weighting 48 – 49 grams or less (i.e., the weight of a star-nosed mole or smaller). This is well above the body weights estimated for some early therians, whose size has been compared to those of modern mice or shrews. For example, the early eutherian *Juramaia* is estimated to have weighted around 15-17 grams (Luo et al., 2011), for which we predict a dominant frequency of around 14.3 – 15.0 kHz.

A second possibility is that mammals’ high-frequency hearing and vocalizations, were adopted for navigation early in their evolutionary history. As mentioned above, early mammals probably vocalized at high-frequencies due to their small size, and high-frequency sounds benefit from improved directionality at the expense of poorer sound propagation (Wiley & Richards 1978; Surlykke et al., 2008). Directional vocalizations and hearing are important requirements for biosonar systems used in navigation, and the need for more directional calls is in part responsible for the high-frequencies of bat echolocation vocalizations (Jakobsen et al., 2013). Echolocation has been considered a form of autocommunication by some authors (Bradbury & Vehrencamp 1998; Jones et al., 2021), and besides bats, it has been confirmed in mice, odontocetes and humans (Schenkman & Nilsson 2010; Panyutina et al., 2017; He et al., 2021; Brinkløv et al., 2023). Some form of echo-guided navigation has been suggested for a tarsier, a shrew, the tenrec and the hippopotamus (Gould 1965; Maust-Mohl et al., 2018; Gursky 2019; Gleason et al., 2023), and its likely to be formally described for other mammals in the future. Overall, there seem to be widespread echolocating potential among mammals, and its development tends to be associated with limited visibility conditions (Brinkløv et al., 2023). Just like most living mammals carry the signature of their nocturnal past in their visual system (Hall et al., 2012), it is possible that the biosonar potential of modern mammals is a remnant of a small, nocturnal and potentially echolocating ancestor. If this were the case, this ancestor was probably a rather unspecialized, but still functional, echolocator. Perhaps with navigation behaviour more similar to soft-furred tree mice (Panyutina et al., 2017; Volodin et al., 2018; He et al., 2021), or even echolocating birds (Brinklov et al., 2013), than the sophisticated sonar systems of bats and odontocetes.

Third, besides its possible role in communication with others or autocommunication, high-frequency hearing may have had a function in prey detection. The songs of katydids have been part of the nocturnal soundscape for at least 165 million years (Gu et al., 2012), and fossil evidence suggests that the hearing systems of mammals has evolved in temporal correspondence with an increase in the frequency content of katydid songs (Xu et al., 2022).

In summary, the ultimate causes underlying the evolution of mammalian hearing are far from clear and remain largely speculative. None of the three possibilities mentioned above are mutually exclusive, and these may not be the only possibilities. Indeed, it is likely that the small size of early mammals and the presence of parent-offspring communication, together with their nocturnal and insectivorous habits all contributed to some degree to the evolution of the definitive therian auditory system, and therefore, permitted their vocal diversification.

## Conclusion

Tetrapods have adopted vocalizations for communication in multiple contexts and virtually every living species produces a distinctive call. Here we have shown that tetrapod vocal evolution has occurred at different tempos and modes, with mammals being outstanding for their faster rates of vocal evolution and higher frequencies. We argue that the outstanding vocal diversity observed in mammals is allowed by their unique auditory systems, which evolved in the context of a small, parental-caring, nocturnal, insectivorous and potentially echolocating ancestor. More generally, we conclude that the evolution of communicative signals cannot be understood independently from the evolution of the perceptual systems responsible for their detection. The notion that particularities in the sensory system of perceivers can promote the evolution of novel signals or the elaboration of pre-existing ones is not new (Ryan et al., 1990; Ryan & Cummings 2013). We interpret the results of our study as an expression of the same phenomenon at a large macroevolutionary scale. The critical role of auditory systems in permitting or limiting the elaboration achievable by vocal systems is not completely surprising, especially considering that signal detection is a *sine qua non* for communication to occur. In a sense, it is fair to say that when it comes to the evolution of vocal systems, the last word is in the ears of the perceivers. As a final thought experiment, we suggest that if birds had evolved a mammalian-like hearing system, they would very probably be the most outstanding vocalizers alive. After all, in its origin the larynx was not devoted to vocal production, but rather to preventing objects from entering the lungs (Negus 1929). As such, the larynx of modern non-avian tetrapods is characterized by this dual vocal and respiratory functionality. In contrast, the specialized syrinx of birds has the single function of being a vocal production organ (Fitch & Hauser 2003), and therefore is free from the trade-offs associated with multi-functionality.

## Author contributions

MIM, MM, and WH conceived the study. MM collected the data. MM and MIM analyzed the data. MIM wrote the manuscript with input from MM, JE and WH.

## Acknowledgements

We are grateful to the members of the Sensory Ecology and Evolution Amsterdam laboratory for the insightful conversations around the evolution of vocalizations. MIM is supported by Becas Chile 2018—CONICYT scholarship (number: 72190501). The authors declare no conflicts of interest.

## Data accessibility statement

All the data will be made available on Dryad upon acceptance.

