## Supplemental Tables for "Tetrapod vocal evolution: higher frequencies and faster rates of evolution in mammalian vocalizations"

**Supplementary Table 1:** Sensitivity analysis to evaluate the influence of random reductions in overall sample size on parameter estimates. We performed 1000 simulations for each category of data reduction (i.e., 10, 20, 30, 40, and 50%), each time randomly removing a different set of species. The average differences between the full model (i.e., including all the species) and the reduced one (i.e., with a sub-sample of the species) is expressed in percentage and as the raw difference.

| **Parameter** | **% of species removed** | **% of significant parameters** | **Mean change (%)** | **Mean difference** |
|---|---|---|---|---|
| Intercept | 10 | 100 | 0.44 | 0.100 |
| | 20 | 100 | 0.70 | 0.213 |
| | 30 | 100 | 0.86 | 0.404 |
| | 40 | 100 | 1.13 | 0.567 |
| | 50 | 100 | 1.27 | 0.736 |
| Slope | 10 | 100 | 2.12 | -0.115 |
| | 20 | 100 | 3.28 | -0.216 |
| | 30 | 100 | 4.37 | -0.426 |
| | 40 | 100 | 5.85 | -0.558 |
| | 50 | 100 | 7.14 | -0.790 |

**Supplementary table 2:** Sensitivity analysis to evaluate the influence of major tetrapod clades on the tetrapod weight-frequency allometry. All the species from each major lineage (mammals, birds or frogs) were completely removed, and new PGLS slopes and intercepts were computed from this reduced dataset. The estimates from this reduced model were compared against the parameters estimated from the full model (i.e., including all clades), and differences expressed as raw differences and percentage of change. The P-value corresponds to the significance of the test that evaluates differences between these estimates. Additionally, we computed a mean null estimate obtained from a random sample of tetrapods. This random sample of mammals, birds and frogs was simulated 1000 times, and the number of species randomly chosen was the same as species in the clade removed, effectively controlling for differences in the sampling size between groups (mammals are underrepresented in the sample relative to birds and frogs)

| | | | | | | | | |
|---|---|---|---|---|---|---|---|---|
| Clade removed | Number of species | Full model slope | Reduced model slope | Difference in slopes | Change (%) | P-value | Mean null estimate | P-value random. |
| Mammalia | 103 | -0.235 | -0.185 | 0.050 | 21.2 | **1.17e-29** | -0.237 | **0.000** |
| Aves | 404 | -0.235 | -0.275 | -0.040 | 17.0 | **2.01e-35** | -0.244 | **0.029** |
| Amphibia | 366 | -0.235 | -0.245 | -0.011 | 4.5 | **5.03e-23** | -0.242 | 0.430 |
| Clade removed | Number of species | Full model intercept | Reduced model intercept | Difference in intercepts | Change (%) | P-value | Mean null estimate | P-value random. |
| Mammalia | 103 | 3.770 | 3.492 | -0.278 | 7.4 | **1.17e-29** | 3.776 | **0.000** |
| Aves | 404 | 3.770 | 3.880 | 0.110 | 2.9 | **2.01e-35** | 3.797 | 0.052 |
| Amphibia | 366 | 3.770 | 4.040 | 0.270 | 7.2 | **5.03e-23** | 3.794 | **0.000** |

**Supplementary table 3:** Comparison of models testing for differences in the regimes of residual frequency evolution between clades using OU models. A regime is defined by a unique combination of trait optimum ($\theta$, or in this case value at the root, $a_0$), strength of pull towards the optimum ($\alpha$), and evolutionary rate ($\sigma^2$). The different models assume different scenarios of evolution between mammals, birds and frogs, going from all regimes equal to all regimes different.

| Model regimes | Model description | Degrees of freedom | AIC | ΔAIC |
|---|---|---|---|---|
| mammal ≠ bird = frog | Common regime for birds and frogs | 7 | -4.319 | 0 |
| mammal ≠ bird ≠ frog | All regimes different | 9 | -1.386 | 2.933 |
| mammal = bird ≠ frog | Common regime for mammals and birds | 7 | 136.974 | 141.292 |
| mammal = frog ≠ bird | Common regime for mammals and frogs | 5 | 140.803 | 145.122 |
| mammal = bird = frog | All regimes equal | 4 | 181.169 | 185.488 |

**Supplementary table 4:** Result from the best-fit model to study differences in the regimes of residual frequency evolution between mammals, birds and frogs. The best-fit model (see **Supp. Table 3**) included a common-regime for birds and frogs, and a different regime for mammals.

| | | | | | | | | | |
|---|---|---|---|---|---|---|---|---|---|
| Common-regime OU model: | | | | | | | | | |
| | $\sigma^2$ | $a_{0, mammal}$ | $a_{0, bird}$ | $a_{0, frog}$ | $\alpha$ | k | log(lik) | | |
| Value | 1.059 | 0.453 | 0.0008 | -0.303 | 0.298 | 5 | -85.58 | | |
| S.E. | 0.112 | 0.235 | 0.118 | 0.155 | 1.587 | | | | |
| Multi-regime OU model: | | | | | | | | | |
| | $\sigma^2_{mammal}$ | $\sigma^2_{bird} = \sigma^2_{frog}$ | $a_{0,mammal}$ | $a_{0, bird}$ | $a_{0, frog}$ | $\alpha_{mammal}$ | $\alpha_{bird} = \alpha_{frog}$ | k | log(lik) |
| Value | 4.352 | 0.709 | 0.464 | 0.001 | -0.302 | 2.548 | 0.507 | 7 | 9.16 |
| S.E. | 1.515 | 0.086 | 0.358 | 0.095 | 0.123 | 4.838 | 1.671 | | |
| Likelihood ratio: 189.49; $P < 0.0001$ (based on $\chi^2$) | | | | | | | | | |

$\sigma^2$: evolutionary rate.

$a_0$: estimated ancestral estate at the root of each lineage, also known as phylogenetic mean.

$\alpha$: pull strength towards the optimum.

k = degrees of freedom.
